# Parallel molecular mechanisms underlie convergent evolution of the exaggerated snout phenotype in East African cichlids

**DOI:** 10.1101/2022.01.13.476207

**Authors:** Anna Duenser, Pooja Singh, Laurène Alicia Lecaudey, Christian Sturmbauer, Craig Albertson, Wolfgang Gessl, Ehsan Pashay Ahi

**Author notes:** Authors of Correspondence: Ehsan Pashay Ahi, Christian Sturmbauer.

## Abstract

Studying instances of convergent evolution of novel phenotypes can shed light on the evolutionary constraints that shape morphological diversity. Cichlid fishes from the East African Great Lakes are a prime model to investigate convergent adaptations. However, most studies on cichlid craniofacial morphologies have primarily considered bony structures, while soft tissue adaptations have been less intensely scrutinised. A rare example of an exaggerated soft tissue phenotype is the formation of a snout flap. This tissue flap develops from the upper lip and has evolved in only one cichlid genus from Lake Malawi and one genus from Lake Tanganyika. To investigate the molecular basis of snout flap convergence, we used mRNA sequencing to compare two species with snout flap (*Labeotropheus trewavasae* and *Ophthalmotilapia nasuta*) to their close relatives without snout flaps (*Tropheops tropheops* and *Ophthalmotilapia ventralis*) from Lake Tanganyika and Malawi. Our analysis revealed a greater complexity of differential gene expression patterns underlying the snout flap in the younger adaptive radiation of Lake Malawi than in the older Lake Tanganyika radiation. We identified 201 genes that were repeatedly differentially expressed between species with and without the snout flap in both lakes, suggesting that the pathway that gives rise to snout flaps is evolutionarily constrained, even though the flaps play very different functions in each species. The convergently expressed genes are involved in proline and hydroxyproline metabolism, which have been linked to human skin and facial deformities. Additionally, we also found enrichment for transcription factor binding sites upstream of differentially expressed genes such as members of the FOX transcription factor family, especially *foxf1* and *foxa2*, which also showed an increased expression in the flapped snout and are linked to nose morphogenesis in mammals, as well as *ap4* (tfap4), a transcription factor showing reduced expression in the flapped snout with an unknown role in the development of craniofacial soft tissues. As genes involved in cichlids snout flap development are associated with many human mid-line facial dysmorphologies, our findings imply a conservation of genes involved in mid-line patterning across vastly distant evolutionary lineages of vertebrates.

**Significance statement:** The study of the evolution of similar physical traits across taxa can give insight into the molecular architecture underlying shared phenotypes. This has mostly been studied in bony structures, while soft tissue traits have been less intensely covered. We investigated the exaggerated snout in cichlid species from Lake Malawi and Lake Tanganyika and found that many genes involved in the development of the snout flap are also associated with mid-line dysmorphologies in humans, implying a conservation across distant vertebrate lineages.

## Introduction

Convergent evolution of phenotypes, reflecting particular ecological specializations, is a ubiquitous characteristic of adaptive radiations (Schluter & Nagel 1995; Losos et al. 1998; Rundle et al. 2000; Rüber et al. 1999). Cichlid adaptive radiations from the East African Great lakes display an impressive array of repeated morphological traits (Kocher et al. 1993), including a few dramatic examples such as nuchal humps and hypertrophied lips (Machado-Schiaffino et al. 2014; Manousaki et al. 2013; Colombo et al. 2013; Baumgarten et al. 2015; Lecaudey et al. 2019, 2021). However, the evolution of such phenotypic novelties is not well understood, but a comparative approach can shed light on the genetic and developmental mechanisms that reconfigure the body plan and give rise to complex traits. Cases of repeated evolution of such phenotypic novelties can thus also help us to understand the constraints that shape morphological diversity.

The overgrowth of craniofacial soft tissues in cichlid fishes has presented some intriguing examples of exaggerated phenotypes in various anatomical regions such as lips, frontal head (nuchal hump) and nose snout (or nose flap) (Colombo et al. 2013; Manousaki et al. 2013; Henning et al. 2017; Lecaudey et al. 2019; Concannon & Albertson 2015; Conith et al. 2018). The exaggerated snout flap is a pronounced projection that emanates from a flap of fibrous tissue just above the upper lip. It is a rare morphological innovation that has only evolved in two tribes of cichlid fishes from East Africa, the modern Haplochromines in Lake Malawi and the Ectodini in Lake Tanganyika (Concannon & Albertson 2015). When this snout is sexually monomorphic, it is thought to be a trophic adaptation that improves feeding efficiency (Konings 2007). When the snout is sexually dimorphic, it is hypothesised to be involved in sexual selection (Konings 2007; Concannon & Albertson 2015). The cichlid snout flap has been studied at the molecular level only in the genus *Labeotropheus* from Lake Malawi where it is sexually monomorphic and functions as a trophic adaptation to efficiently leverage algae from rocks (Concannon & Albertson 2015; Conith et al. 2018). A similar snout structure has also been described in two species from the Ectodini tribe (*Ophthalmotilapia nasuta* and *Asprotilapia leptura*) from Lake Tanganyika. In *A. leptura* it is sexually monomorphic and likely involved in increased foraging efficiency (similar to *Labeotropheus*), whereas in *O. nasuta* it is only found in mature males and is likely a secondary sexual character (Hanssens et al. 1999; Conith et al. 2019). Thus, the exaggerated snout is a convergent phenotype that evolved independently in two cichlid lineages that diverged > 9 MYA (Irisarri et al. 2018; Conith et al. 2019).

In *Labeotropheus*, the snout is evident histologically by the time the yolk is absorbed and exogenous feeding occurs (~1 month post-fertilization) (Conith et al. 2018; Concannon & Albertson 2015), and early formation and growth of the snout is linked to the transforming growth factor beta (TGFβ) signalling pathway (Conith et al. 2018). However, it remains unclear whether (1) similar molecular players are involved in maintaining this phenotype at later life-history stages and if (2) the molecular mechanism of snout formation is the same in other cichlid species that possess a snout. Furthermore, while previous research focused on the TGFβ signalling pathway, a more extensive molecular interaction map of the formation and maintenance of this exaggerated phenotype remains to be unravelled. A transcriptome-wide overview is particularly important since it is well-known that there is molecular cross-talk between the TGFβ signalling pathway and several other pathways which all play a pivotal role in craniofacial morphogenesis and adaptive evolutionary divergence in teleost fishes (Ahi 2016).

In this study, we set out to investigate the molecular mechanisms that underlie the convergent evolution of the exaggerated snout phenotype, in two non-sister cichlid lineages from lakes Tanganyika and Malawi (Figure 1A). We compared two species that develop the snout; (1) *Labeotropheus trewavasae* (tribe Haplochromini) from Lake Malawi and (2) *Ophthalmotilapia nasuta* (tribe Ectodini) from Lake Tanganyika (Figure 1). As controls, we used two closely related species within each tribe that do not develop such a structure; (1) the Lake Malawi mbuna species *Tropheops tropheops* (Haplochromini) and (2) the Lake Tanganyika featherfin cichlid *Ophthalmotilapia ventralis* (Ectodini) (Figure 1). We used mRNA-sequencing to quantify gene expression differences between the exaggerated snout and non-snout tissues for each lake. Altogether, we identified parallel and non-parallel molecular mechanisms that underlie the evolution of the snout flap in Lake Malawi and Lake Tanganyika cichlids. Our study design provides valuable information on convergent regulatory mechanisms underlying the morphogenesis of a unique hypertrophic facial soft tissue in cichlids, which also exhibit striking similarity to those mechanisms driving craniofacial development and mid-line patterning in other vertebrates including humans.

**Fig. 1.**
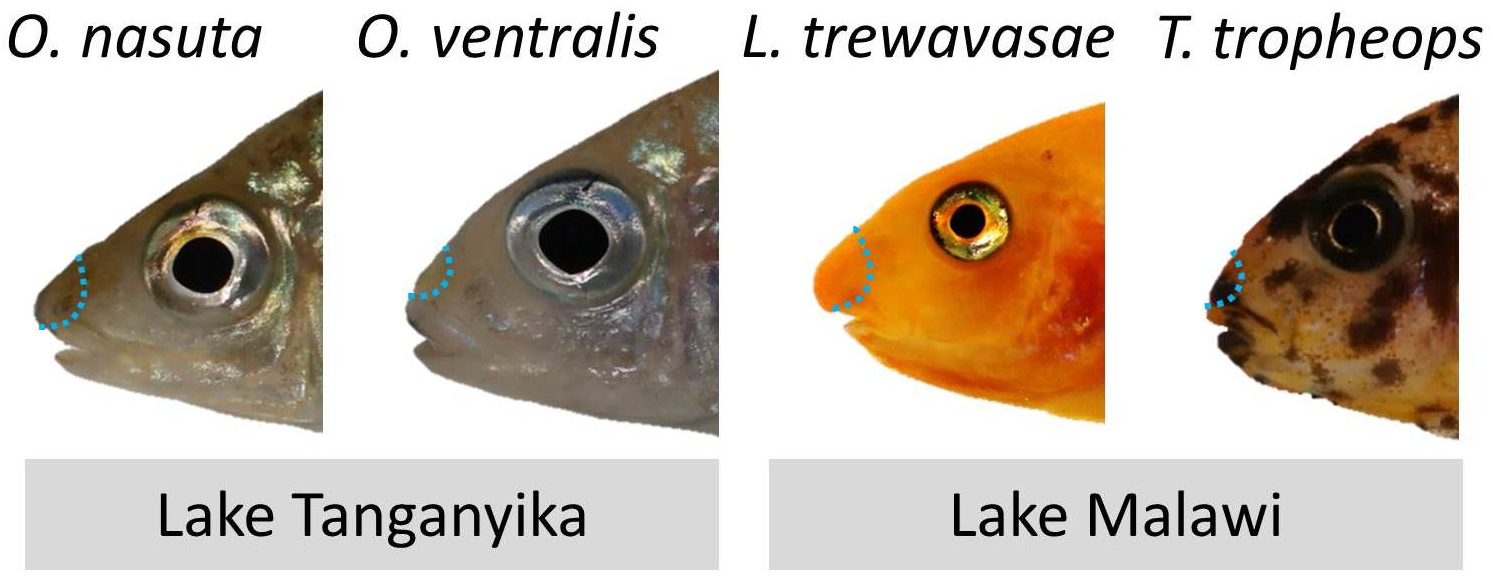
Convergent cases of snout evolution. East African cichlid species used in this study. The area of the soft tissue that was dissected is depicted by blue dashed lines.

## Results

### RNA-sequencing, gene expression and downstream analyses

The RNA-sequencing resulted in between 6.7 and 15.8 million reads per sample and after filtering of low-quality reads, between 4.6 and 11.1 million reads were retained for each sample (Supplementary File S1). The raw data of sequence reads have been deposited in the Sequencing Read Archive (SRA) of NCBI (accession number: PRJNA770252). The pairwise comparisons between species of each lake radiation resulted in 832 differentially expressed genes for the comparison of *O. nasuta* versus *O. ventralis*, while the comparison between *L. trawavasae* versus *T. tropheops* yielded 4292 differentially expressed genes. Between these both results we identified an overlapping list of 201 differentially expressed (DE) genes which were distinct between the flapped snout versus the non-flapped snout regions in both lakes (Figure 2A) (Supplementary File S2). Among the DE genes, 74 genes showed upregulation and 96 genes showed downregulation in snout tissues in both comparisons, whereas 31 genes showed expression differences in opposite directions across the comparisons for each Lake (Figure 2B-D). The heatmap clustering of the DE genes showed that there are at least two major branches in each group of up- or down-regulated gene sets, while the clustering of the DE gene with opposite expression pattern also revealed the presence of two major branches (Figure 2B-D). These clustering structures indicate distinct transcriptional regulations within each group which potentially originated from the effects of different upstream regulators.

**Fig. 2.**
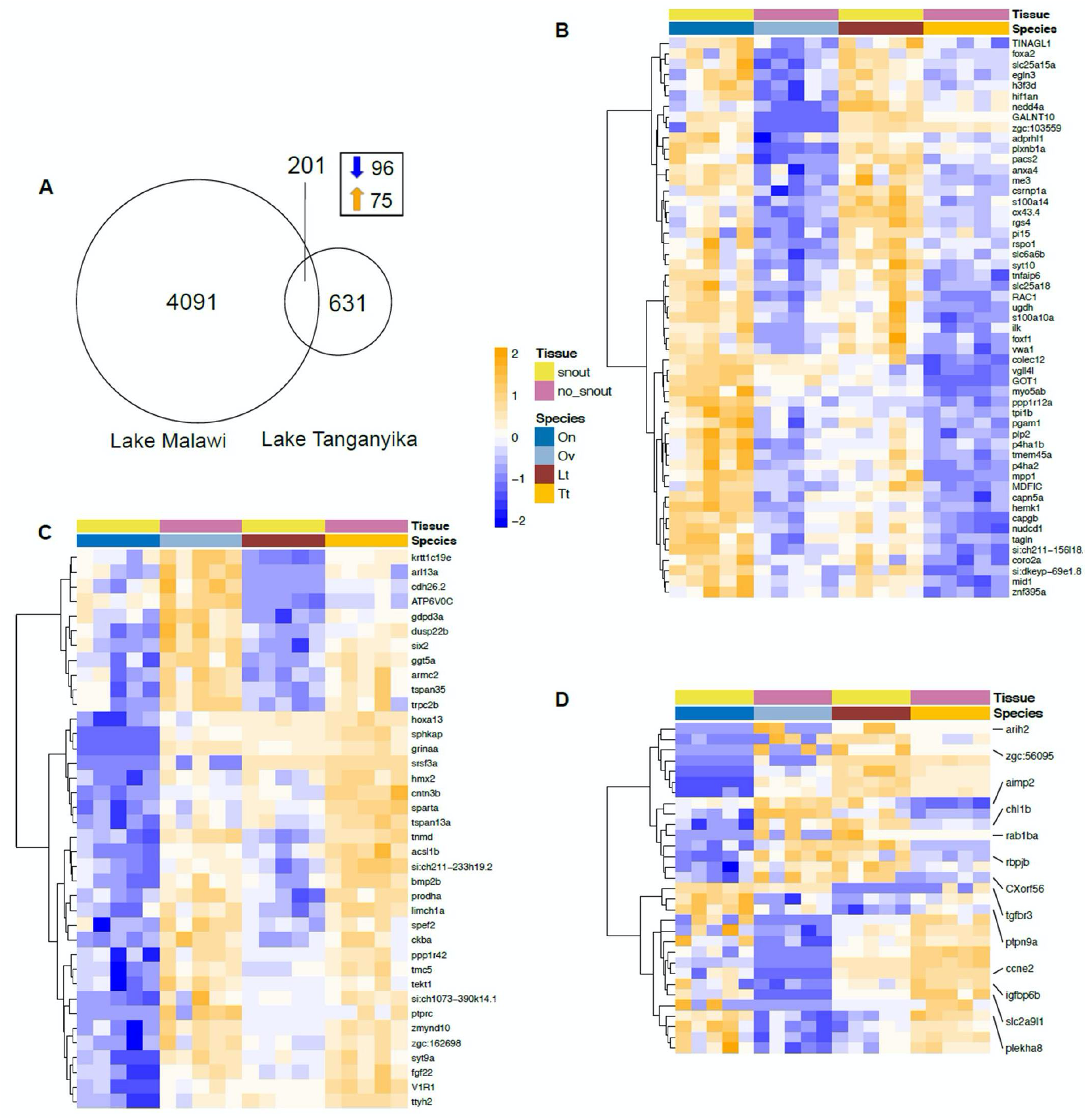
Differentially expressed genes in the snout regions. (**A**) Venn diagram of 201 genes with differential expression between the snout regions (“snout” and “no snout”) which overlap between the two comparisons. Dendrogram clusters of the overlapping annotated genes showing upregulation (**B**), and downregulation (**C**) in expression in the flapped snout tissue, as well as those showing differential expression in both comparisons but in opposite direction (**D**). Orange and blue shadings indicate higher and lower relative expression respectively. *Ophthalmotilapia ventralis* (Ov)*, Ophthalmotilapia nasuta* (On)*, Labeotropheus trewavasae* (Lt), *Tropheops tropheops* (Tt).

We next performed gene ontology enrichment analysis using the list of 201 DE genes as the input, and the result showed significant enrichment of GO terms for several biological processes such as amino acid metabolism (particularly proline related metabolic processes), Wnt and Notch signalling pathways, and regulation of cell adhesion (Figure 3A).

**Fig. 3.**
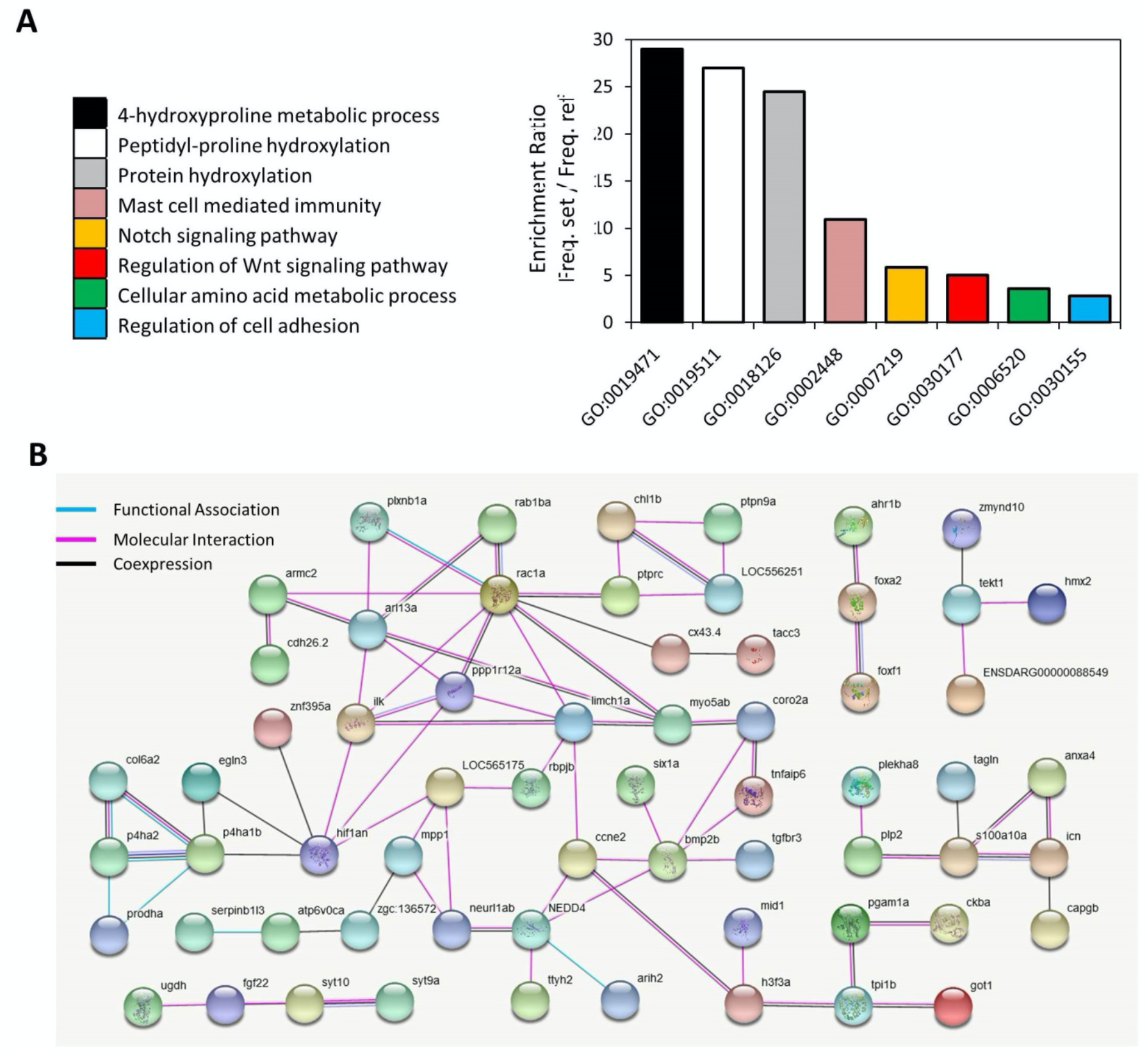
Functional analyses of the overlapping differentially expressed genes in flapped snout. (**A**) Enrichment for gene ontologies of biological processes using the shared 201 differentially expressed genes and Manteia online tool. (**B**) Functional interactions between the differentially expressed genes predicted based on zebrafish databases in STRING v10 (http://string-db.org/).

We also applied the same list of DE genes for interactome analysis which demonstrated a large, interconnected network of genes with molecular and functional associations. Some of the genes in the network formed an interaction hub with the highest level of associations with other genes such as *bmp2b, hif1an* and *rac1a*, suggesting their more pivotal role in the formation of the flapped snout structure in cichlids (Figure 3B). Furthermore, we conducted TF binding motif overrepresentation analysis on the upstream regulatory sequences of the DE genes through MEME tool (Bailey et al. 2009). In total, seven motifs were enriched on the upstream regulatory sequences of at least 40 out of 201 DE genes (Table 1). Next, we checked the similarities of the enriched motifs with known TF binding sites in vertebrates and at least 11 TF candidates were identified to potentially bind to those motifs.

**Table 1.**
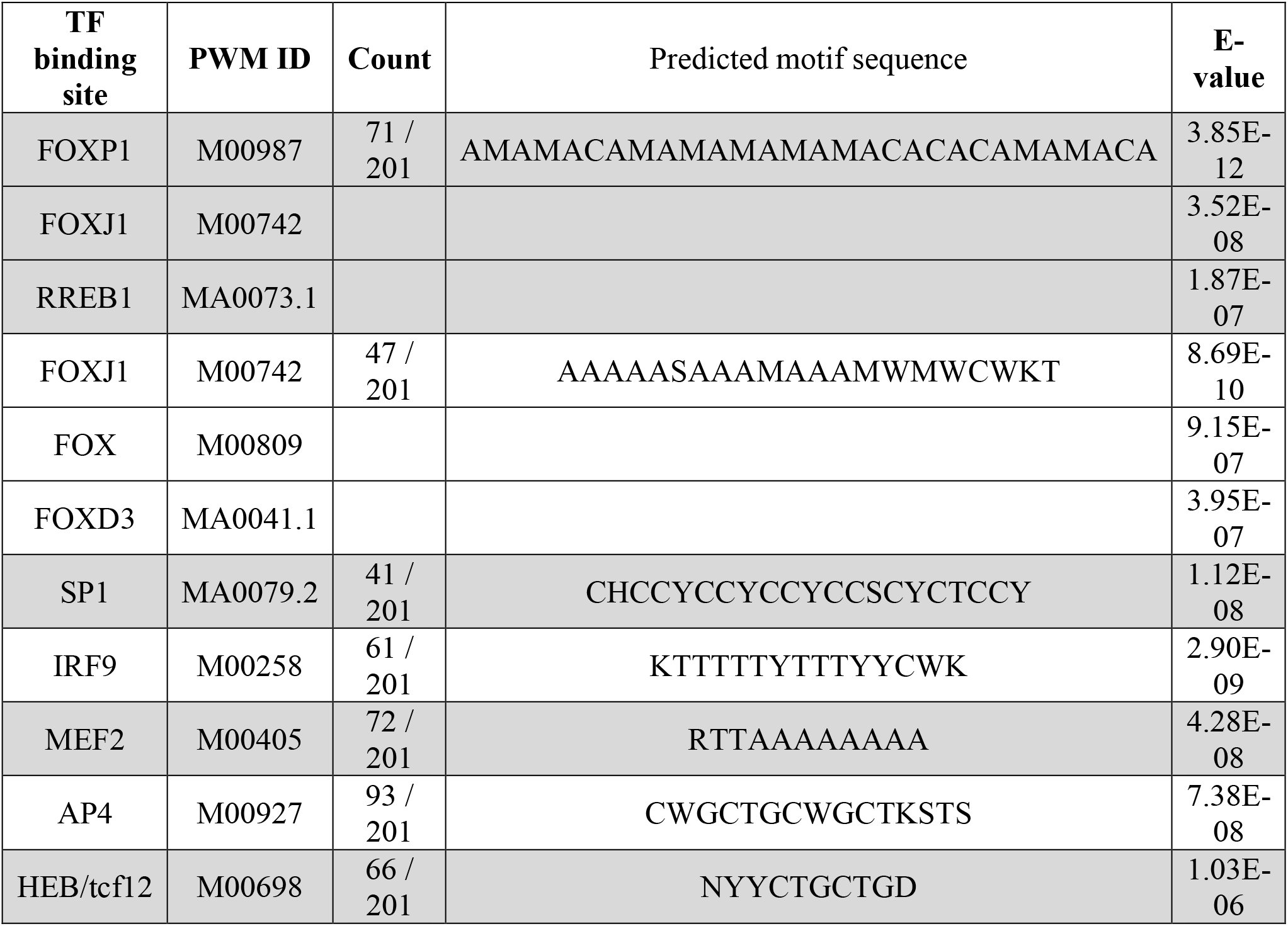
Predicted motifs and upstream regulators potentially binding to them. Enriched motifs on upstream regulatory sequences of the DE genes are presented in degenerated sequence format. PWD IDs indicate positional weight matrix ID of predicted binding sites and E-values refer to matching similarity between the predicted motif sequences and the PWD IDs. The count implies the number of genes containing the predicted motif sequence on their regulatory region.

### Expression analysis by qPCR

Validation of DE genes from RNA-seq was accomplished via quantitative RT-PCR (qPCR), normalised to stably expressed reference genes (Kubista et al. 2006). In our previous studies of East African cichlids, we found that validation of reference gene(s) is an essential step for every species and tissues (Ehsan P. Ahi et al. 2020; Ehsan Pashay Ahi et al. 2020; Pashay Ahi & Sefc 2018; Ahi et al. 2019). We chose six candidate reference genes with a small log2 fold change and the lowest coefficient of variation (CV) throughout all the samples (Supplementary File S2). Based on the rankings by the three software tools, BestKeeper, geNorm and NormFinder, only one of the candidate reference genes, *pak2b*, showed consistent stability, i.e. always ranked among top two most stable reference genes (Table 2). Thus, we used the Cq value of *pak2b* in each sample to normalize the relative gene expression levels of our target genes.

**Table 2.**
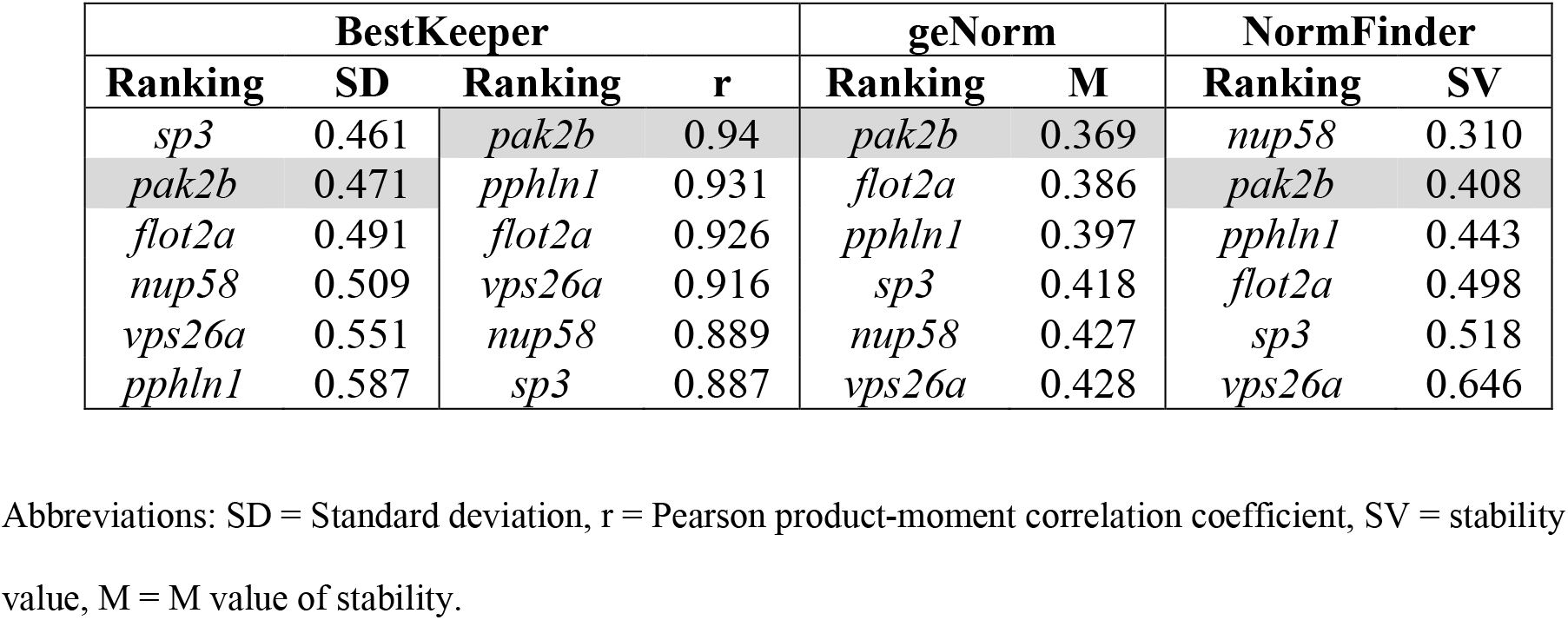
Ranking of reference genes in the nose tissue samples using three different algorithms.

| BestKeeper |  |  |  | geNorm |  | NormFinder |  |
| --- | --- | --- | --- | --- | --- | --- | --- |
| Ranking | SD | Ranking | r | Ranking | M | Ranking | SV |
| *sp3* | 0.461 | *pak2b* | 0.94 | *pak2b* | 0.369 | *nup58* | 0.310 |
| *pak2b* | 0.471 | *pphln1* | 0.931 | *flot2a* | 0.386 | *pak2b* | 0.408 |
| *flot2a* | 0.491 | *flot2a* | 0.926 | *pphln1* | 0.397 | *pphln1* | 0.443 |
| *nup58* | 0.509 | *vps26a* | 0.916 | *sp3* | 0.418 | *flot2a* | 0.498 |
| *vps26a* | 0.551 | *nup58* | 0.889 | *nup58* | 0.427 | *sp3* | 0.518 |
| *pphln1* | 0.587 | *sp3* | 0.887 | *vps26a* | 0.428 | *vps26a* | 0.646 |

Abbreviations: SD = Standard deviation, r = Pearson product-moment correlation coefficient, SV = stability value, M = M value of stability.

Among the DE genes identified by RNA-seq, we chose 12 genes with a known role in nose morphogenesis and/or other related functions in craniofacial development mainly based on genetic studies in humans (Table 3), together with eight predicted upstream TFs (including *ap4, foxd3, foxj1, foxp1, irf9, mef2a, rreb1a* and *sp1*) for qPCR analysis (Figure 4).

**Fig. 4.**
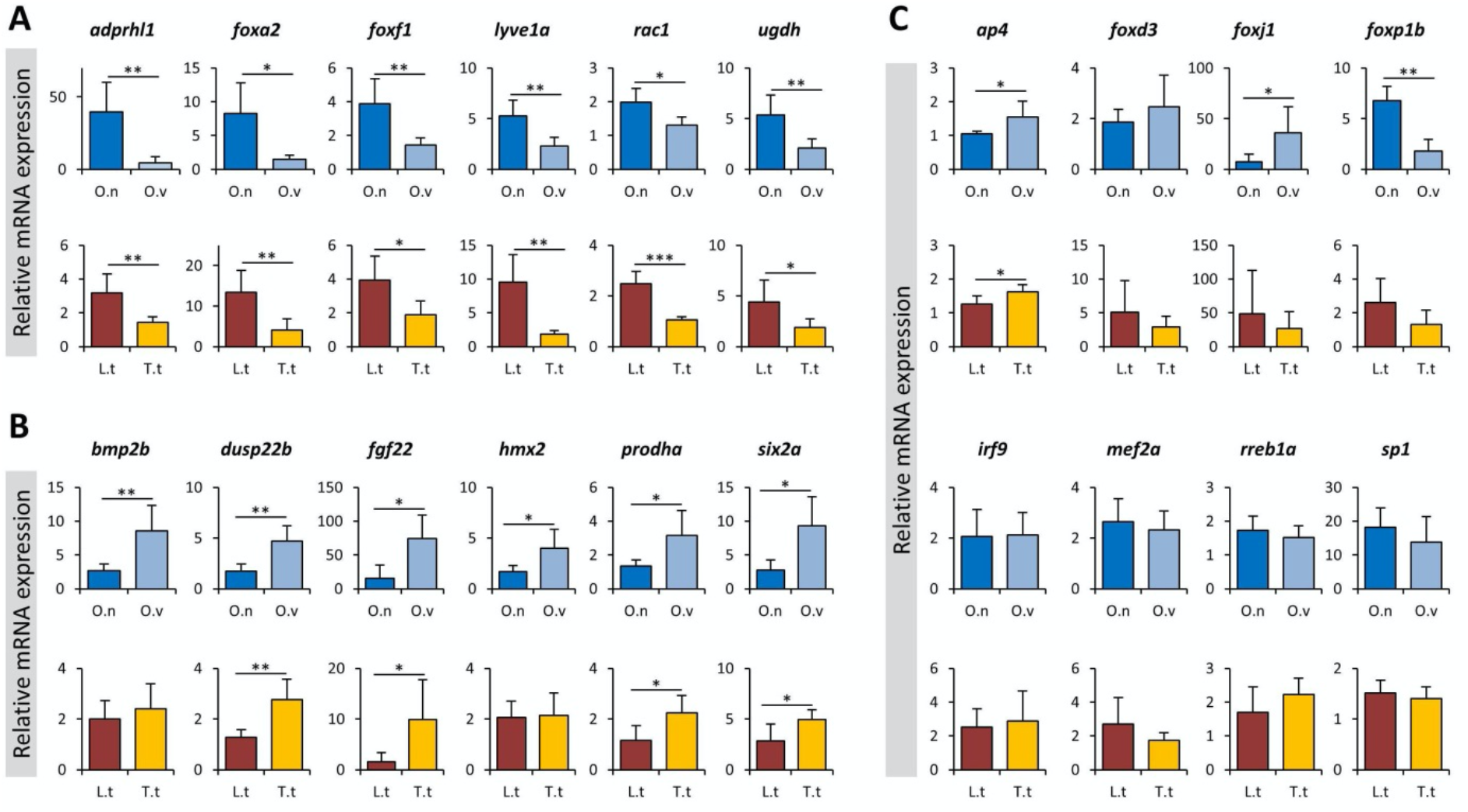
qPCR expression analysis of a selected set of candidate genes. The bars indicate mean and standard deviation of RQ expression values for five biological replicates per species. The asterisks above the bar represent significant expression differences (*P < 0.05; **P < 0.01; ***P < 0.001). *Ophthalmotilapia ventralis* (O.v), *Ophthalmotilapia nasuta* (O.n)*, Labeotropheus trewavasae* (L.t), *Tropheops tropheops* (T.t)

**Table 3.**
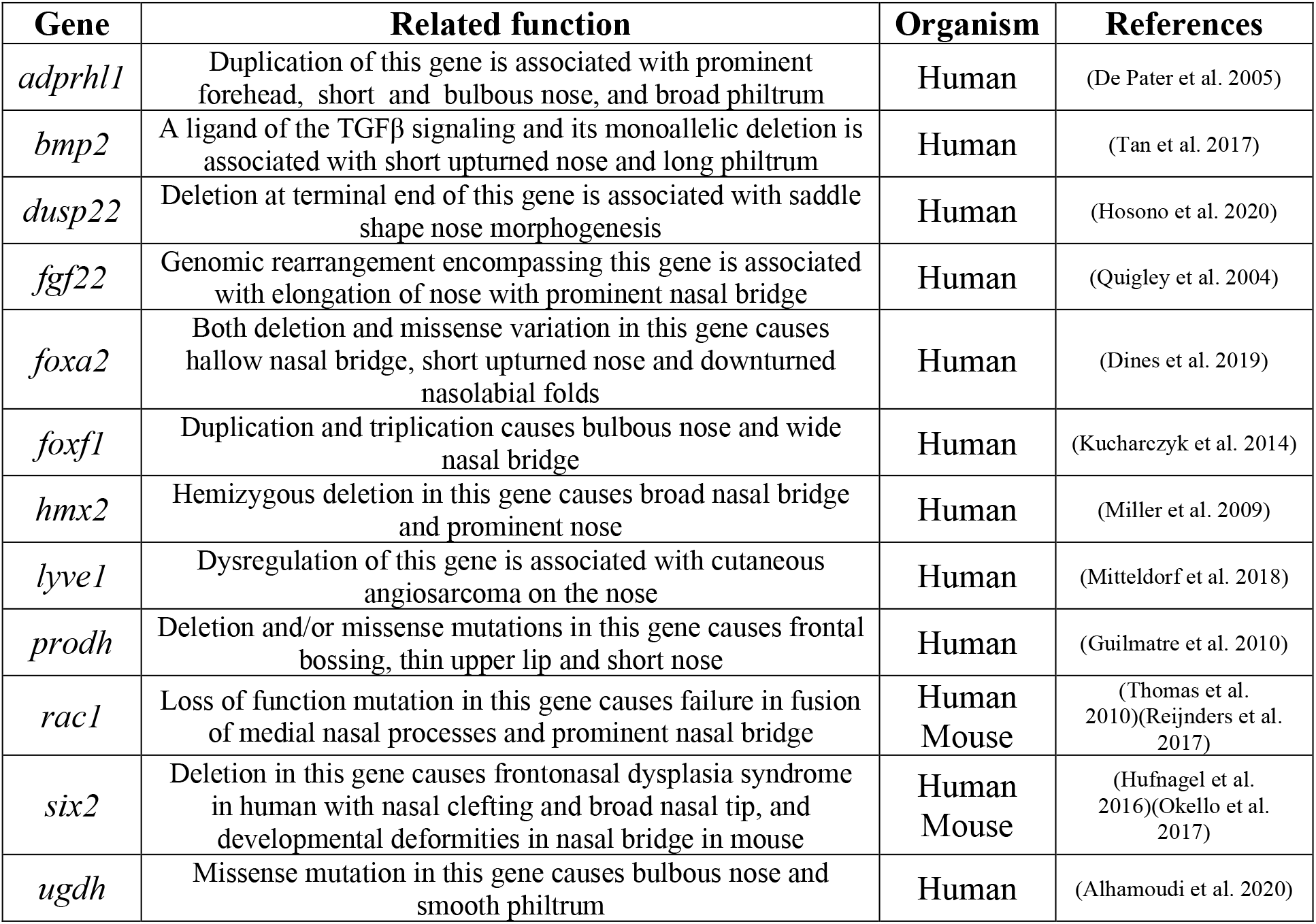
A selected set of differentially expressed genes in the flapped snout regions of studied cichlids with known related functions in nose morphogenesis in mammalian models.

| Gene | Related function | Organism | References |
| --- | --- | --- | --- |
| *adprhl1* | Duplication of this gene is associated with prominent forehead, short and bulbous nose, and broad philtrum | Human | (De Pater et al. 2005) |
| *bmp2* | A ligand of the TGFβ signaling and its monoallelic deletion is associated with short upturned nose and long philtrum | Human | (Tan et al. 2017) |
| *dusp22* | Deletion at terminal end of this gene is associated with saddle shape nose morphogenesis | Human | (Hosono et al. 2020) |
| *fgf22* | Genomic rearrangement encompassing this gene is associated with elongation of nose with prominent nasal bridge | Human | (Quigley et al. 2004) |
| *foxa2* | Both deletion and missense variation in this gene causes hallow nasal bridge, short upturned nose and downturned nasolabial folds | Human | (Dines et al. 2019) |
| *foxf1* | Duplication and triplication causes bulbous nose and wide nasal bridge | Human | (Kucharczyk et al. 2014) |
| *hmx2* | Hemizygous deletion in this gene causes broad nasal bridge and prominent nose | Human | (Miller et al. 2009) |
| *lyve1* | Dysregulation of this gene is associated with cutaneous angiosarcoma on the nose | Human | (Mitteldorf et al. 2018) |
| *prodh* | Deletion and/or missense mutations in this gene causes frontal bossing, thin upper lip and short nose | Human | (Guilmatre et al. 2010) |
| *rac1* | Loss of function mutation in this gene causes failure in fusion of medial nasal processes and prominent nasal bridge | Human<br>Mouse | (Thomas et al. 2010)(Reijnders et al. 2017) |
| *six2* | Deletion in this gene causes frontonasal dysplasia syndrome in human with nasal clefting and broad nasal tip, and developmental deformities in nasal bridge in mouse | Human<br>Mouse | (Hufnagel et al. 2016)(Okello et al. 2017) |
| *ugdh* | Missense mutation in this gene causes bulbous nose and smooth philtrum | Human | (Alhamoudi et al. 2020) |

Based on the RNA-seq results, six of these candidate genes displayed up-regulation in expression in the flapped snout (*adprhl1*, *foxa2*, *foxf1*, *lyve1a*, *rac1* and *ugdh*), while the six other candidate genes (*bmp2b*, *dusp22b*, *fgf22*, *hmx2*, *prodha* and *six2a*) showed a down-regulation in expression in the flapped snout. The results of qPCR analysis confirmed that almost all of the genes showed expression patterns similar to RNA-seq results, except for *bmp2b* and *hmx2* which showed no significant difference between the snout regions of L.t and T.t. Among the predicted TFs only *ap4* showed consistent differences across both comparisons displaying a slightly reduced expression in both species with protruded snouts (O.n and L.t). This indicates potential transcriptional repressor effects of *ap4* on the downstream genes in the hypertrophic snout region. Two members of FOX transcription factors, *foxj1* and *foxp1*, also showed expression differences but only in one of the comparisons (O.n vs O.v). Altogether, the qPCR results demonstrate consistency between RNA-seq and qPCR results confirming the validity of our transcriptome data analysis.

## Discussion

Biologists have been fascianted with divergent as well as convergent morphological evolution as it contributes to our understanding of the interplay between ecological opportunity and genetic constraints. The snout flap of *L. trevawasae* is thought to have evolved under natural selection (Concannon & Albertson 2015), as it plays a distinct role in the foraging efficiency for algal scraping (Konings 2007; Conith et al. 2019). Additionally, no difference in snout flap size has been detected between male and female of *Labeotropheus* and its formation has been shown to coincide with the time point of independent foraging, further supporting its function (Concannon & Albertson 2015). In contrast, only *O. nasuta* males show the distinct snout flap, implicating its role in mate choice (Concannon & Albertson 2015). Furthermore, both sexes of *O. nasuta* are planktivorous suction feeders, a feeding adaptation that is presumably not enhanced by a snout flap, although the snout of males continues to grow with increasing age (Hanssens et al. 1999). In a comparison of tissue types of the snout flap it has been found that the snout of *Labeotropheus fuelleborni* contains a lot more of intermaxillary ligament and loose connective tissue (80%) than the snout of *O. nasuta* (50%) (Conith et al. 2019). The morphological convergence of the snout flap across two cichlid radiations allows us to investigate if convergence can also be found at the transcriptional level, even if the morphologies probably possess different functions and differ in tissue composition and life-history.

We found many of DE genes, both upregulated and downregulated, that are associated with craniofacial development and involved in human dysmorphologies, many with mid-line facial defects including those that effect the nose in literature. Among the upregulated genes with related functions were *adprhl1* (De Pater et al. 2005), *angptl2* (Ehret et al. 2015), *colec12* (Zlotina et al. 2016), *cx43* (McLachlan et al. 2005), *foxa2* (Dines et al. 2019), *foxf1* (Kucharczyk et al. 2014), *galnt10* (Starkovich et al. 2016), *got1* (Tomkins et al. 1983), *lyve1* (Mitteldorf et al. 2018), *mdfic* (Kosho et al. 2008), *mid1* (Preiksaitiene et al. 2015)(Hüning et al. 2013), *nudcd1* (Selenti et al. 2015), *pacs2* (Holder et al. 2012), *plxnb1* (Haldeman-Englert et al. 2009), *rac1* (Thomas et al. 2010)(Reijnders et al. 2017), *rspo1* (Wieacker & Volleth 2007), *s100a10* (Sawyer et al. 2007), *slc25a18* (Chen et al. 2013), *slc6a6* (Kariminejad et al. 2015), *ugdh* (Alhamoudi et al. 2020), *vgll4* (Czeschik et al. 2014)(Barrionuevo et al. 2014), and *vwa1* (Giannikou et al. 2012). Among the downregulated genes we also found the following candidates to have such roles; *acsl1* (Yakut et al. 2015), *adgb* (Alazami et al. 2016), *arl13* (Brugmann et al. 2010), *ATP6v0c* (Mucha et al. 2019; Tinker et al. 2021), *bmp2* (Tan et al. 2017), *cntn3* (Ţuţulan-Cuniţa et al. 2012), *dusp22* (Hosono et al. 2020)(Martinez-Glez et al. 2007), *fgf22* (Quigley et al. 2004), *gdpd3* (Dell’Edera et al. 2018), *grina* (Bonaglia et al. 2005), *hmx2* (Miller et al. 2009), *hoxa13* (Fryssira et al. 2011), *il23r* (Rivera-Pedroza et al. 2017), *ppp1r42* (Mordaunt et al. 2015), *prodh* (Guilmatre et al. 2010), *six2* (Hufnagel et al. 2016)(Okello et al. 2017), *srsf3* (Pillai et al. 2019), *syt9* (Sofos et al. 2012), and *trpc2* (Sansone et al. 2014)(Zhang et al. 2010). Interestingly, one of the downregulated genes, *pi15*, is known as an important molecular player in beak formation in birds (Nimmagadda et al. 2015). Even among the overlapping DE genes which showed opposing expression patterns between the two comparisons, we still found at least four genes to have been associated with craniofacial mid-line defects in other vertebrates, including *ccne2* (Jain et al. 2010), *plekha8* (Schulz et al. 2008), *rab1b* (Alwadei et al. 2016), *rbpj* (Nakayama et al. 2014) and *tgfbr3* (Lopes et al. 2019). These findings demonstrate that similar sets of genes are involved in mid-line patterning and growth across evoulutionary distant vertebrates. Thus, functional studies investigating their specific role in divergent morphogenesis of snout structures in fish can provide valuable information about the conserved molecular mechanisms underlying the formation of facial soft tissues (Powder & Albertson 2016).

Conducting gene ontology enrichment analysis on the list of DE genes also revealed the involvement of several biological processes such as proline and hydroxyproline metabolisms, regulation of cell adhesion, as well as Notch and Wnt signalling pathways in the formation of the flapped snout in these cichlid species. Interestingly, a defective proline and hydroxyproline metabolisms has been already associated with a range of skin and facial deformities including abnormal nose morphogenesis in humans (Kiratli & Satilmiş 1998; Kretz et al. 2011; Zaki et al. 2016; Baumgartner et al. 2016). A defective proline metabolism is known to severely affect collagen formation and extracellular matrix integrity, and subsequently cell adhesion (Velez et al. 2019; Karna et al. 2020; Xinjie et al. 2001; Javitt et al. 2019; Noguchi et al. 2020). The lack of proline hydroxylation in collagen by reduced prolyl 4-hydroxylase activity can directly affect integrin binding and cell adhesion mechanisms (Sipila et al. 2018). Interestingly, we found genes involved in 4-hydroxyproline metabolic process as the most enriched biological process, which suggests changes in proline hydroxylation as top candidate of metabolic changes during the formation of exaggerated snout in cichlids. In addition, it has been recently shown that the biosynthesis of proline is tightly regulated by transforming growth factor-beta (Tgfβ) (Schwörer et al. 2020), a transcription factor that is also playing an important role in the early development of the flapped snout structure in cichlids (Conith et al. 2018). Although, we did not detect differential expression of *Tgfβ1* itself, components of this pathway were identified (e.g., *tgfbr3*), and both of the enriched signalling pathways, Wnt and Notch, have evolutionary conserved crosstalk with Tgfβ mediated signal in regulation of various molecular, cellular and developmental events (Attisano & Labbé 2004; Chesnutt et al. 2004; Arnold et al. 2019; Klüppel & Wrana 2005; Ahi 2016). In addition, both Wnt and Notch signalling pathways are known to play a pivotal role in craniofacial development and morphogenesis including the formation of middle structures including nasal structures (Penton et al. 2012; Pakvasa et al. 2020; Brugmann et al. 2007; Wang et al. 2011).

A proposed model for Tgfβ and Notch crosstalk suggests that induction of Tgfβ signalling is required for the early establishment of cell-cell contacts in different tissues, whereas later induction of Notch signal stabilizes the Tgfβ mediated effects (Klüppel & Wrana 2005). This allows the cells to react to a new environment through induction of alternative genetic programs controlling differentiation or migration (Klüppel & Wrana 2005). In the context of the snout, it is possible that activation of Tgfβ is required for early snout induction (Conith et al. 2018) and that continued snout growth is maintained via Notch signalling. This potential time dependent crosstalk may be mediated through the downstream targets of Notch and Tgfβ signals, since it is shown that both signals can regulate similar target genes (Klüppel & Wrana 2005; De Jong et al. 2004), including *foxa2*, a member of the FOX family of transcription factors (both signals suppress *foxa2* expression) (Kondratyeva et al. 2016; Liu et al. 2012). In our study, we found upregulation of *foxa2* in the flapped snout region, and interestingly, a recent study in human shows that a deletion in *Foxa2* can cause specific facial deformities including a shallow nasal bridge, a short upturned nose, and a downturned nasolabial fold (Dines et al. 2019). In addition, we found *rbpjb*, a major transcription factor mediating canonical Notch signal in various cell types (Tanigaki et al. 2002) to be differentially expressed in the flapped snout of both species. *rbpj* is shown to directly regulate a receptor of Tgfβ signal (*Tgfbr1*) in mice, thus making a reciprocal positive regulatory loop between the two pathways (Valdez et al. 2012). We also found another receptor of Tgfβ signal (*tgfbr3*) to show a similar expression pattern as *rbpjb* raising the possibility of the existence of such a reciprocal regulatory loop in flapped snout cichlids. In human, a deletion in *Rbpj* gene has been linked to developmental defects in brain and abnormal thickening of the nose and lip (Nakayama et al. 2014). Another genome-wide study revealed that *Rbpj* acts as a direct upstream regulator of Wnt signalling in mammalian stem cells (Li et al. 2012). On the other hand, Bmp2 signal which is regarded as a molecular cross point between Smad/Tgfβ and Notch pathways (De Jong et al. 2004), mediates its signal through *Tgfbr3* (Hill et al. 2012). It is already known that Bmp2 can regulate Notch signal and its downstream target genes (De Jong et al. 2004). Previous studies in cichlids had proposed variations in Bmp expression as a molecular player in adaptive morphological divergence in different skeletal structures (Gunter et al. 2013; Albertson et al. 2005; Ahi et al. 2017; Hulsey et al. 2016). We found down-regulation of *bmp2b* expression suggesting that a key regulator linking both pathways is affected in the flapped snout region of the cichlid species in this study. Furthermore, deletion of *Bmp2* in human has been reported to cause a range of facial deformities including shortened nose, anteverted nares, elongation of the philtrum and changes in the thickness of lips (Tan et al. 2017). Taken together, these findings suggest potentially complex interactions between Notch and Tgfβ signals in the formation and possibly the maintenance of the flapped snout structure in cichlids.

Finally, we also conducted enrichment for TF binding sites on regulatory sequences of DEGs and found several potential binding sites for TFs that may play a role in the formation of a flapped snout. The most represented TF binding sites belonged to members of FOX transcription factor family, e.g. *foxd3, foxj1* and *foxp1*. In previous studies of East African cichlids, both *foxd3* and *foxp1* were found to act upstream of a gene network involved in exaggerated fin elongation (Pashay Ahi & Sefc 2018; Ahi et al. 2019). Additionally, *foxp1* was recently identified as a key upstream regulator of genes involved in the formation of the hypertrophic lip in another East African cichlid species from Lake Tanganyika (Lecaudey et al. 2021). None of the predicted FOX members (*foxd3*, *foxj1* and *foxp1*) displayed consistent differential expression across both comparisons, but interestingly, the consensus binding site for FOX transcription factor family was also among the predicted binding sites. It is, therefore, possible that the two other FOX members identified by RNA-seq and qPCR, *foxf1* and *foxa2*, are the key regulators of the entire list of DEGs, since they might bind to the consensus FOX binding site. In addition, both *foxf1* and *foxa2* displayed consistently increased expression in the flapped snout in both comparisons, and are also implicated in the nose morphogenesis in mammals (Dines et al. 2019; Kucharczyk et al. 2014). We have recently found *foxf1* among the regulators of lip hypertrophy in an East African cichlid species as well (Lecaudey et al. 2021), suggesting a potentially important and general role of *foxf1* in soft tissues exaggeration in cichlids, and given the potential role of *foxd3* and *foxp1* in cichlid fin exaggeration, FOX genes may play a more general role in tissue elaboration in vertebrates. Among the other predicted TF binding site we found overrepresentation of binding motif for *tcf12*, a transcription factor with known roles in cranio-skeletal development; particularly in the morphogenesis of the frontal bone and cranial vault thickening in mammals (Piard et al. 2015; Sharma et al. 2013). Moreover, we have previously identified *tcf12* as a potential key player in the formation of a nuchal hump in an East African cichlid (Lecaudey et al. 2019). In this study, however, we did not detect its expression in the snout region by differential gene expression or qPCR analysis. The only predicted TF with consistent expression difference in both comparisons was *ap4* (or *tfap4*), i.e. showing slight but significant reduced expression in the flapped snout. *ap4* encodes a member of the basic helix-loop-helix-zipper (bHLH-ZIP) family, and can act as a transcriptional activator or repressor on a variety of downstream target genes mediating cell fate decisions (Wong et al. 2021). The exact role of *ap4* in craniofacial morphogenesis of soft tissues is unclear, however, deletions in a genomic region containing this gene appeared to cause facial dysmorphisms in human such as prominent beaked nose and micrognathia (Gervasini et al. 2007). Future functional studies are required to verify the role of *ap4* in formation and morphogenesis of craniofacial soft tissues in fish.

## Conclusions

The snout flap in *Labeotropheus trewavasae* and *Ophthalmotilapia nasuta* is a striking and rare example of an exaggerated soft tissue trait that has evolved repeatedly in the cichlid radiations of Lake Malawi and Lake Tanganyika, albeit for different functions. Comparing the transcriptional landscape of the snout flap tissue of these two species with the snout of close relatives that do not develop such a structure, we identified 201 genes that were repeatedly recruited to give rise to the snout flap phenotype even after > 9 MYA of divergence. Our study provides support for a change in proline hydroxylation, a mechanism also linked to human facial deformations, to be a mechanism for metabolic changes involved in the formation of the snout flap in fish. Additionally, we found indications of complex interactions between the transforming growth factor-beta (Tgfβ), regulating the biosynthesis of proline, and Notch signalling, associated with morphogenesis and craniofacial development, in the formation and maintenance of the snout flap. Upstream of the differentially expressed genes we identified transcription factors belonging to the FOX family (especially *foxf1* and *foxa2*) which are both linked to the morphogenesis of the nose in mammals and *ap4* a transcription factor that showed reduced expression in the species with snout flap, but with an unknown role in craniofacial soft tissue formation. We want to emphasise that the identification of genes involved in snout morphogenesis in fish can shed light on the conserved molecular mechanisms crucial for the development and shaping of facial soft tissue. In the future it would be important to build on these findings and confirm the reuse of these genes and pathways across more distant teleost groups.

## Methods

### Fish rearing and tissue sampling

Five captive bred males of each *O. nasuta*, *O. ventralis*, and five captive bred females of *L. trewavasae* and *T. tropheops* were raised and kept in a large tank (approximately 450 litres) containing multiple stony shelters to decrease competition stress. All specimens were at the young adult stage and have been fed with the same diet, Tropical multi-ingredient flakes suitable for omnivorous cichlids. The two species in each comparison were sampled at the same time when the protrusion of the flapped snout had already appeared (Figure 1). To perform the dissections, we used a solution with 0.3 g MS222 per 1L water to euthanize the fish, and similar snout regions, an area above the upper lip encompassing the nostrils which includes epidermis, dermis and the underlying soft connective tissues, were sampled for each fish (Figure 1). The sampled snout tissues for each individual were placed into separate tubes containing RNAlater (Qiagen) and stored at −20 C°. The sacrificing of fish followed the guidelines of the Federal Ministry of Science, Research and Economy of Austria according to the regulations of the BMWFW.

### RNA extraction and cDNA synthesis

Total RNA was extracted from 20 dissected snout tissue samples (5 biological replicates per species) following the TRIzol method (Thermo Fischer Scientific). Each dissected sample included epidermis, dermis, and the underlying fibrous/connective tissues of the specified nose regions (Figure 1). Tissue samples were placed into tubes containing 1 ml of TRIzol with a ceramic bead (1.4mm) and homogenized using a FastPrep-24 Instrument (MP Biomedicals, CA, USA). RNA extraction followed the protocol of TRIzol RNA extraction from Thermo Fischer Scientific. A DNA removal step with DNase followed the extraction (invitrogen). The total RNAs were dissolved in 50 μl nuclease-free water and their concentrations were quantified through a Nanophotometer (IMPLEN GmbH, Munich, Germany). We measured the quality of RNAs with the R6K ScreenTape System using an Agilent 2200 TapeStation (Agilent Technologies) and RNA integrity numbers (RIN) above 7 were aimed at for all samples. To synthesize cDNA for qPCR analysis, we used 500 ng of the total RNA per sample and followed the manufacturer’s protocol of the High Capacity cDNA Reverse Transcription kit (Applied Biosystems), and the resulted cDNAs were diluted 1:4 to be used for the qPCR reaction.

### RNA-seq library preparation and gene expression quantification

To attain transcriptome data of the snout tissues, we conducted RNA-seq library preparation with 1000 ng of total RNA per tissue sample as input and following the protocol of the Standard TruSeq Stranded mRNA Sample Prep Kit (Illumina) with indexing adapters. The library qualities were assessed using D1000 ScreenTape and reagents (Agilent) on a TapeStation 2200 machine (Agilent). In order to reach an optimal quantity recommended for sequencing, we diluted the libraries and pooled them with equal molar concentration for each library. The RNA-sequencing was conducted in the NGS Facility at Vienna Biocenter Core Facilities (VBCF, Austria) on an Illumina HiSeq2500 and generated between 6.7 and 15.8 million paired-end reads with 125bp length per sample (Supplementary File S1). Raw reads were de-multiplexed based on unique barcodes by the same facility. The quality of the reads was assessed with FastQC (v0.11.8) (Andrews 2012), and reads were filtered for a quality > 28 and a minimum length of 70 bp with Trimmomatic (v0.3.9) (Bolger et al. 2014). Reads were aligned to the *O. niloticus* reference genome (Conte et al. 2017) of the University of Maryland using RNAstar (v2.7.3.a) (Dobin et al. 2013). To check the mapping statistics, we used samtools idxstats (v1.9) (Danecek et al. 2021) and further merged the single files for species and Lake with picard (v2.21.7) (Picard Toolkit. 2019. Broad Institute, GitHub Repository. https://broadinstitute.github.io/picard/). We used StringTie (v.2.0.6) (Pertea et al. 2015) to assemble the alignments into potential transcripts without a reference. This step was conducted separately for single files (per biological replicate) and the merged files (per species and per Lake). The single files per biological replicate were further merged into species. This process of repeated merging steps was implemented to reduce the probability of false positives. To assess the accuracy of the mapping we used gffcompare (v0.11.2) (Pertea & Pertea 2020) to compare our annotations to the reference annotation. Subsequently we filtered for monoexonic transcripts that were not contained in our reference and the transcripts assigned the class code ‘possible polymerase run-on’ by gffcompare. As the maximum intron length of the *O. niloticus* reference is 200000 bp, we also filtered for that in the produced annotation. The expression estimates for each transcript were based on these annotations and generated with StringTie (v.2.0.6) with no multimapping allowed. The final raw count matrices were produced from the expression estimates with a Perl script from the griffith lab (https://github.com/griffithlab/rnaseq_tutorial/blob/master/scripts/stringtie_expression_matrix.pl) and the code used in this analysis is available at this github repository (https://github.com/annaduenser/snout_flap_RNAseq).

Differential expression analysis was conducted using DESeq2 (Love et al. 2014) in R (R Core Team 2017) running comparisons for each Lake separately. DESeq2 estimates variance-mean dependence based on a model using negative binomial distribution using the raw counts (Love et al. 2014). A false discovery rate of p < 0.05 was chosen as the cutoff. For the downstream analysis, an enrichment step for gene ontology (GO) terms of biological processes was conducted using the intersection list of DE genes between the Lakes and Manteia, a free online tool for data mining of vertebrate genes (Tassy & Pourquié 2014). In addition, we investigated the functional interactions between the products of DE genes through STRING v10 (http://string-db.org/), a knowledge based interaction prediction tool, and zebrafish databases for protein interactomes (Szklarczyk et al. 2017).

### Primer design and qPCR

We designed the qPCR primers on conserved regions of the selected candidate genes by aligning their assembled sequences to their already available homologous mRNA sequences from *Ophthalmotilapia ventralis* (Böhne et al. 2014), *Metriaclima zebra, Pundamilia nyererei, Neolamprologus brichardi* and *Astatotilapia burtoni* (Brawand et al. 2014), as well as *Oreochromis niloticus*. After aligning the conserved sequence regions across the abovementioned East African cichlids, we identified the exon junctions (using CLC Genomic Workbench, CLC Bio, Denmark, and annotated genome of *Astatotilapia burtoni* in the Ensembl database, http://www.ensembl.org). The primer designing steps were conducted as described previously (Pashay Ahi & Sefc 2018; Ahi et al. 2019) using Primer Express 3.0 (Applied Biosystems, CA, USA) (Supplementary file S3). The qPCR was performed based on the protocol provided by Maxima SYBR Green/ROX qPCR Master Mix (2X) (Thermo Fisher Scientific, Germany) following the guidelines for optimal experimental set-up for each qPCR run (Hellemans et al. 2007). The qPCR program was set for 2 min at 50°C, 10 min at 95°C, 40 cycles of 15 sec at 95°C and 1 min at 60°C, followed by an additional step of dissociation at 60°C – 95°C. The primer efficiency (E values) for each gene was calculated through standard curves generated by serial dilutions of pooled cDNA samples. The standard curves were run in triplicates and calculated using the following formula: E = 10[−1/slope] (Supplementary file S3).

In order to select stably expressed candidate reference genes, we filtered for genes with a low log2 fold change and subsequently ranked the remaining genes according to low coefficient of variation. The top six most stable genes shared across the transcriptome comparisons were selected as candidate reference genes. After qPCR expression analysis of the six genes across all samples, we ranked them based on their expression stability by three different algorithms: BestKeeper (Pfaffl et al. 2004), NormFinder (Andersen et al. 2004) and geNorm (Vandesompele et al. 2002). We used the Cq values of the top most stable reference genes to normalize Cq values of target genes in each sample (ΔCq target = Cq target – Cq reference). The relative expression levels (RQ) were calculated by 2^-ΔΔCq^ method (Pfaffl 2001) and the log-transformed RQ values were used for t-tests to calculate the statistical differences.

## Acknowledgements

The authors thank Holger Zimmermann and Stephan Koblmüller for sharing their precious knowledge on cichlid fishes of Lake Tanganyika, Sylvia Schäffer for sharing her experience on RNA-seq library preparation, and Martin Grube and his lab for technical assistance and access to their real-time PCR System. The authors acknowledge the financial support by the University of Graz.

## Author Contributions

EPA, CA and CS conceived the project. WG contributed to fish husbandry and photography, and EPA and AD conducted the sampling and tissue dissection. AD, EPA and LL conducted the RNA lab work. AD, PS, LL, EPA contributed to the analyses and all authors to manuscript writing. CS and EPA contributed to funding. This work was supported by the Austrian Science Fund [project number P29838] awarded to CS. All authors approved the final version of the manuscript.

## Competing financial interests

The authors declare no competing interests.

## Ethical approval

No experiments were conducted on the fish prior to sampling, so an ethics approval is not required according to the Austrian animal welfare law. Fish keeping and sacrifice was carried out in our certified aquarium facility in accordance with the Austrian animal welfare law.

## Data availability

The data underlying this article are available in the Sequencing Read Archive (SRA) of NCBI at https://www.ncbi.nlm.nih.gov/ and can be accessed with PRJNA770252.

## Notes

### Competing Interest Statement

The authors have declared no competing interest.

